# The thymic microenvironment gradually modulates the phenotype of thymus-homing peripheral conventional dendritic cells

**DOI:** 10.1101/2021.08.27.457792

**Authors:** Susanne Herppich, Michael Beckstette, Jochen Huehn

## Abstract

Thymic conventional dendritic cells (t-DCs) are crucial for the development of T cells. A substantial fraction of t-DCs originates extrathymically and migrates to the thymus. Here, these cells contribute to key processes of central tolerance like the clonal deletion of self-reactive thymocytes and the generation of regulatory T (Treg) cells. So far, it is only incompletely understood which impact the thymic microenvironment has on thymus-homing conventional DCs (cDCs), which phenotypic changes occur after the entry of peripheral cDCs into the thymus and which functional properties these modulated cells acquire. In the present study, we mimicked the thymus-homing of peripheral cDCs by introducing *ex vivo* isolated splenic cDCs (sp-DCs) into re-aggregated thymic organ cultures (RTOCs). Already after two days of culture, the transcriptomic profile of sp-DCs was modulated and had acquired certain key signatures of t-DCs. The regulated genes included immunomodulatory cytokines and chemokines as well as co-stimulatory molecules. After four days of culture, sp-DCs appeared to have at least partially acquired the peculiar Treg cell-inducing capacity characteristic of t-DCs. Taken together, our findings indicate that peripheral cDCs possess a high degree of plasticity enabling them to quickly adapt to the thymus-specific microenvironment. We further provide indirect evidence that thymus-specific properties such as the efficient induction of Treg cells under homeostatic conditions can be partially transferred to thymus-homing peripheral cDC subsets.

## Introduction

The thymic microenvironment is formed by a coordinated set of cellular components as well as various soluble proteins establishing a complex network of interactions. Importantly, within this multilayered network each component is closely connected and interdependent.^1^ Among the cellular components are antigen-presenting cells (APCs), including thymic epithelial cells (TECs) and thymic conventional dendritic cells (t-DCs). These have an extraordinary impact on the development of T cells and on the establishment of a functional and highly diverse T cell receptor (TCR) repertoire.^2^ Within the cortex and medulla of the thymus, T cell lineage commitment as well as a cascade of discrete consecutive differentiation steps occur. These steps, which lead to the generation of T cells bearing a virtually unlimited number of diverse TCRs, facilitate protection against a vast spectrum of pathogens. However, elimination of potentially autoreactive clones is required to ensure tolerance towards innocuous and self-antigens.^3^ Thus, self-reactive thymocytes are either driven to apoptotic cell death by negative selection or differentiate into regulatory T (Treg) cells, which are characterized by the expression of the transcription factor Foxp3 and essential for the maintenance of immune homeostasis and self-tolerance.^2-4^

Conventional DCs (cDCs) within the thymus play a critical role in the establishment of central tolerance. The compartment of t-DCs is comprised of approximately 70 % cDC1s (CD8α^+^SIRPα^-^ t-DCs) and 30 % cDC2s (CD8α^-^SIRPα^+^ t-DCs).^5^ Thymic cDC1s, also referred to as resident t-DCs, arise primarily within the thymus from an early, yet undefined thymic progenitor. Due to their XCR1 expression, CD8α^+^SIRPα^-^ t-DCs co-localize with medullary TECs (mTECs), expressing the XCR1 ligand XCL1.^6-8^ Thereby, resident CD8α^+^SIRPα^-^ t-DCs are able to cross-present tissue-restricted antigens that are expressed in mTECs.^9^ In contrast, thymic cDC2s (CD8α^-^SIRPα^+^ t-DCs) are migratory t-DCs that originate extrathymically and migrate from the periphery to the thymus. Within the thymus, CD8α^-^SIRPα^+^ t-DCs are enriched at the cortico-medullary junction as well as around small vessels.^10-12^ These migratory t-DCs can capture and display blood-born antigens, but also present peripheral antigens.^13^

Accumulating evidence suggests that both resident and migratory t-DCs play a role in negative selection and also contribute to the thymic Treg cell development in a non-redundant manner.^2, 4, 14, 15^ Yet, while CD8α^-^SIRPα^+^ migratory t-DCs were shown to harbor a superior capacity to induce Treg cells *in vitro*, splenic conventional DCs (sp-DCs) possess only poor Treg cell induction capacity when compared to bulk t-DCs.^9, 13, 16-18^ This raises the question whether the thymic microenvironment has the capacity to modulate the phenotype of thymus-homing peripheral cDCs and instruct them with efficient Treg cell-inducing properties.

In the present study, we mimicked the thymus-homing of peripheral cDCs by introducing *ex vivo* isolated sp-DCs into re-aggregated thymic organ cultures (RTOCs). This technique provides the possibility to investigate *in vitro* the interplay of the complex thymic microenvironment with any cell population of interest.^19^ We found that the thymic microenvironment has the capacity to rapidly shift the phenotype of sp-DCs towards a t-DC phenotype on transcriptome level. Only a short residence within the RTOC likely improved the Treg cell-inducing capacity of sp-DCs, suggesting that the thymic microenvironment harbors a dominant capacity to modulate the functional properties of thymus-homing peripheral cDCs.

## Results

### RTOCs mature over time and sucessfully support thymocyte and TEC development

In order to use RTOCs as a tool to study the impact of the thymic microenvironment on thymus-homing peripheral cDCs, we first had to ensure that RTOCs roughly mimic the overall cellular composition of a thymus under steady-state conditions. For this purpose, we examined by flow cytometry (see **Figure S1** in Supplementary Material) the frequencies of major thymocyte and TEC populations within RTOCs harvested on day 2 and day 4 of culture, and compared them to *ex vivo* isolated adult thymi. Thymocytes constitute the largest cell population within a thymus. They develop from thymus-seeding progenitors via different CD4^-^CD8^-^ double-negative (DN) stages into CD4^+^CD8^+^ double-positive (DP) thymocytes, which then further differentiate into either CD4 single-positive (SP) or CD8SP cells.^20^ While RTOCs harvested on day 2 possessed only a low proportion of CD4SP and CD8SP thymocytes, which differs significantly from the thymocyte subset composition in *ex vivo* isolated adult thymi, frequencies of SP thymocytes increased during two further days of culture, reaching levels similar to those found in adult thymi (**Figure 1A+B**). Within the CD4SP thymocyte population, the frequency of CD25^+^Foxp3^hCD2+^ Treg cells was significantly reduced in RTOCs when compared to *ex vivo* isolated adult thymi, yet also showing a slight increase over time (**Figure 1A+B**). Similarly, the frequency of CD25^-^ Foxp3^hCD2+^ Treg cell precursors (Foxp3^+^ TregP) also differs significantly between RTOCs harvested on both day 2 and day 4 when compared to *ex vivo* isolated adult thymi. In contrast, the frequency of CD25^+^Foxp3^hCD2-^ Treg cell precursors (CD25^+^ TregP) was strongly decreasing over time with significant differences between RTOCs and *ex vivo* isolated adult thymi observable at day 2 of the culture. Thus, the process of Treg cell maturation within RTOCs seemed to be slower when compared to the maturation of conventional T cells (**Figure 1A+B**). Although the population of DP and DN thymocytes did not follow this overall scheme of maturation, the late DN subpopulations (DN3 and DN4) on day 4 of the culture, separated by the divergent expression of CD25 and CD44, again were closer to frequencies found in adult thymi (**Figure 1A+B**). Viewed as a whole, RTOCs support the development and maturation of major thymocyte populations. Yet, mild alterations between RTOCs harvested at day 4 and adult thymi can still be observed for thymocytes, especially for CD4SP and DN thymocyte subsets.

**Figure 1.**
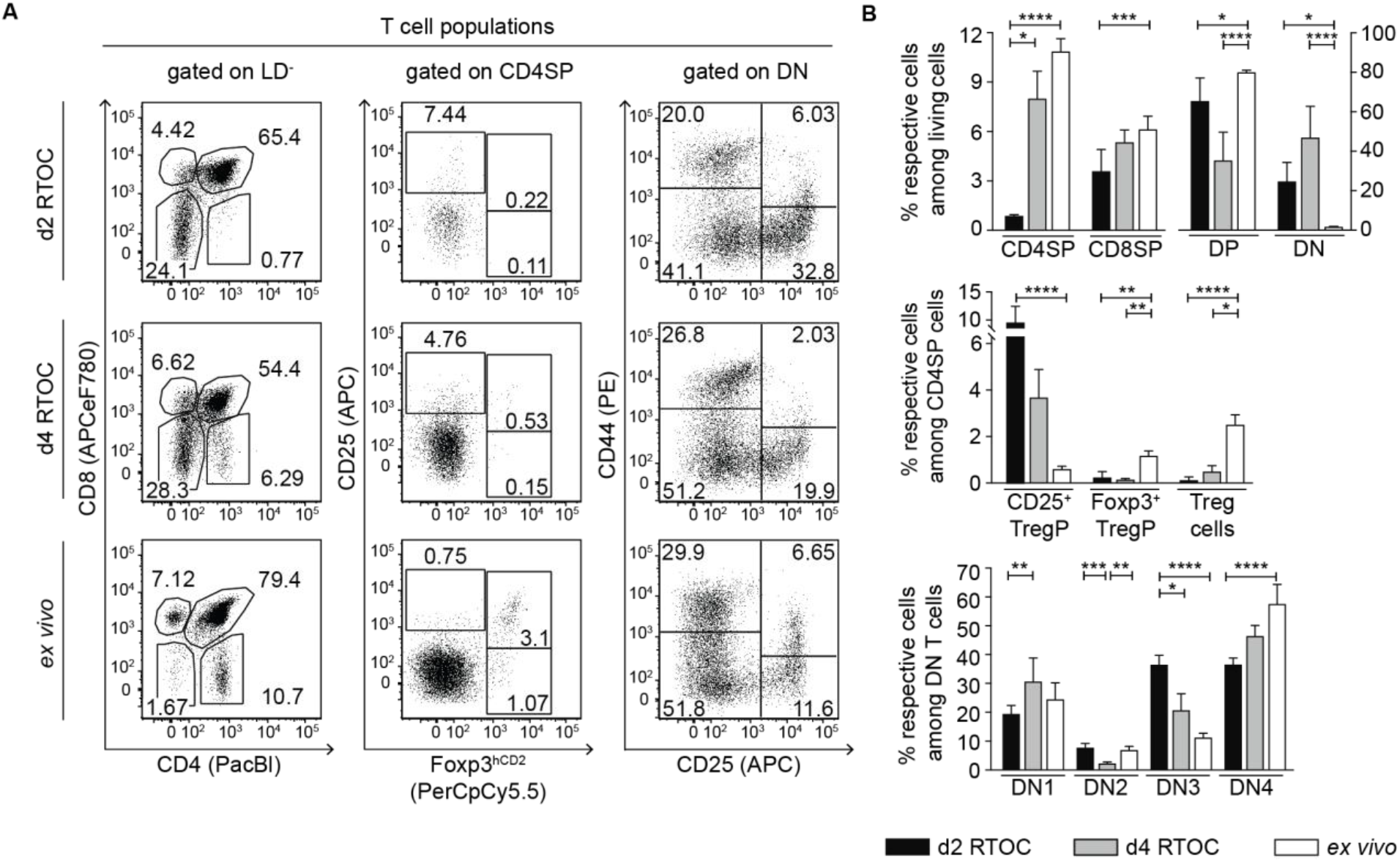
The thymocyte compartment within RTOCs develops over time, approaching population frequencies of adult thymi. RTOCs were generated from single-cell suspensions of pooled thymi isolated from E14.5 to E16.5 fetuses of Foxp3^hCD2^ reporter mice (BALB/c background). After two (d2 RTOCs) or four days (d4 RTOCs), RTOCs were harvested and the thymocyte composition was analyzed by flow cytometry. *Ex vivo* isolated thymi from 4-6 weeks old male Foxp3^hCD2^ reporter mice were analyzed as controls. **(A)** Representative dot plots of thymocyte populations. Numbers indicate the frequencies of cells within the depicted gates. Cells were gated on Live/Dead (LD)-negative cells. **(B)** Graphs show frequencies of indicated thymocyte populations. Data are summarized from two to three independent experiments (mean ± SD; n = 3-5 biological replicates per experiment per group; n = 6-9 biological replicates per group in total). Data of each cell population was independently tested for significance using Kruskal-Wallis test. Significant differences were indicated by * p<0.05; ** p<0.01; *** p<0.001; **** p<0.0001.

TECs are another important cellular component of the thymus. These CD45^-^EpCAM^+^ cells constitute the scaffold of the thymus and strongly interact with thymocytes and t-DCs. It is well-known that the fraction of TECs among total thymic cells decreases early during ontogeny, while the amount of developing thymocytes increases.^1, 21^ This general phenomenon was also observed in the present study as the frequency of CD45^-^EpCAM^+^ TECs among total thymic cells was higher in RTOCs harvested on both day 2 and day 4 when compared to adult thymi (**Figure 2A+B**). TECs consist of two main subsets, medullary and cortical TECs (mTECs and cTECs), named by their localization within the two heterogeneous morphological regions of the thymus – cortex and medulla.^22^ cTECs and mTECs were identified as Ly51^+^UEA-1^-^ and Ly51^-^UEA-1^+^, respectively, among CD45^-^EpCAM^+^ cells.^23^ RTOCs harvested on day 2 and day 4 both showed a high frequency of cTECs and a lower frequency of mTECs among total TECs, while the opposite was observed in *ex vivo* isolated adult thymi (**Figure 2A+B**). Although cTECs tended to decline and mTECs to increase within RTOCs over time, it becomes obvious that these cell populations reached frequencies found in adult thymi noticeably slower when compared to the abovementioned thymocyte populations. Thus, RTOCs harvested on day 4 still showed a reversed ratio of the two TEC populations when compared to *ex vivo* isolated adult thymi. However, both RTOCs harvested on day 2 and day 4 already showed similar frequencies of mTEC subpopulations, as defined by the differential expression of CD80 and MHC II,^24^ when compared to *ex vivo* isolated adult thymi (**Figure 2A+B**). Overall, RTOCs mature over time and possess the capacity to successfully support thymocyte and TEC development. Thus, they are a versatile tool to study diverse aspects of the thymic microenvironment *in vitro*. Yet, it has to be considered that the complete establishment of the TEC compartment takes longer and cannot be completed during the short culture period of the RTOCs.

**Figure 2.**
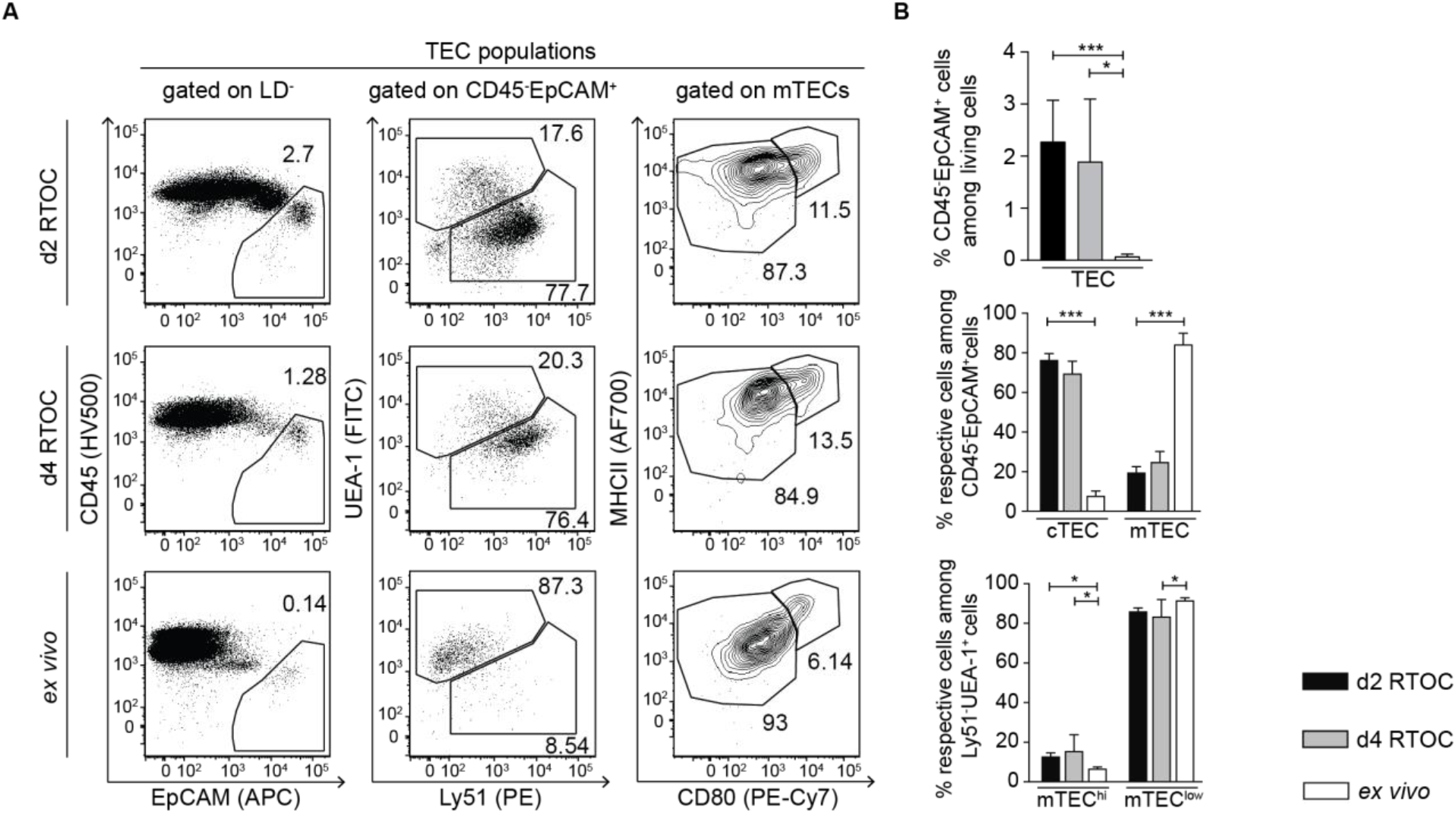
The TEC compartment reaches frequencies found in adult thymi more slowly. RTOCs were generated from single-cell suspensions of pooled thymi isolated from E14.5 to E16.5 fetuses of Foxp3^hCD2^ reporter mice (BALB/c background). After two (d2 RTOCs) or four days (d4 RTOCs), RTOCs were harvested and the TEC composition was analyzed by flow cytometry. *Ex vivo* isolated thymi from 4-6 weeks old male Foxp3^hCD2^ reporter mice were analyzed as controls. **(A)** Representative dot plots for TEC populations. Numbers indicate the frequencies of cells within the depicted gates. Cells were gated on Live/Dead (LD)-negative cells. **(B)** Graphs show frequency of indicated TEC populations. Data are summarized from two to three independent experiments (mean ± SD; n = 3-5 biological replicates per experiment per group; n = 6-9 biological replicates per group in total). Data of each cell population was independently tested for significance using Kruskal-Wallis test. Significant differences were indicated by * p<0.05; ** p<0.01; *** p<0.001.

### Introducing sp-DCs into RTOCs does not negatively affect Treg cell frequencies

To mimic the impact of the thymic microenvironment on thymus-homing peripheral cDCs, we set up RTOCs and introduced CD45^+^Lin^-^CD11c^hi^ sp-DCs *ex vivo* isolated from CD45.1 congenic mice (**Figure S2A-C** in Supplementary Material). RTOCs with corresponding t-DCs were set up as controls. To assess the impact of the introduced sp-DCs on the Treg cell induction within the RTOC, we additionally introduced CTV-labeled CD4SP Foxp3^hCD2-^ thymocytes isolated from adult Foxp3 reporter mice (**Figure S2A+D** in Supplementary Material). In this setting, the RTOCs were harvested and analyzed by flow cytometry after four days of culture to provide suffcient time for an efficient Treg cell induction (**Figure S3** in Supplementary Material). The exogenously added CD45.1^+^ cDCs could be easily distinguished from their endogenous CD45.2^+^ counterparts and made up the majority of the Lin^-^CD11c^hi^ cDC population with a significantly higher frequency when compared to CD45.2^+^ cDCs (**Figure 3A** and **Figure S3A** in Supplementary Material). Likewise, the exogenously added CD4SP thymocytes could be accurately distinguished from endogenous CD4SP thymocytes with the help of the CTV label and made up a slightly, yet significantly higher fraction of the entire CD4SP thymocyte population (**Figure 3B** and **Figure S3B** in Supplementary Material). The flow cytometric analysis revealed that exogenously added t-DCs did not have an impact on the Treg cell-inducing capacity of the RTOCs as similar frequencies of CD25^+^Foxp3^hCD2+^ Treg cells were observed among both the endogenous (CTV^-^) and exogenously added (CTV^+^) CD4SP thymocytes when compared to the RTOCs not receiving additional cDCs (**Figure 3C+D, left**). Importantly, the addition of sp-DCs, which are known to possess a poor Treg cell induction capacity when compared to t-DCs,^16^ did not result in a reduction of the Treg cell frequency in the RTOCs as comparable frequencies of CD25^+^Foxp3^hCD2+^ Treg cells were observed among both endogenous and exogenously added CD4SP thymocytes (**Figure 3C+D, left**). In the same line, similar frequencies of CD25^-^Foxp3^hCD2+^ Treg cell precursors (Foxp3^+^ TregP) were found among both endogenous as well as exogenously added CD4SP thymocytes in RTOCs that either only received CD4SP thymocytes or additionally also t-DCs or sp-DCs (**Figure 3C+D, middle**). In contrast, endogenous CD25^+^Foxp3^hCD2-^ Treg cell precursor (CD25^+^ TregP) were slightly but significantly reduced in RTOCs containing additional t-DCs, while frequencies of CD25^+^ TregP among exogenously added CD4SP thymocytes were mildly yet significantly decreased in RTOCs containing additional sp-DCs (**Figure 3C+D, right**). Furthermore, analysis of the proliferation of the exogenously added CD4SP thymocytes revealed that the addition of t-DCs or sp-DCs to the RTOC had no impact on the proliferation of neither newly generated CD25^+^Foxp3^hCD2+^ Treg cells nor the two Treg cell precursor populations as indicated by comparable geometric mean fluorescence intensities (gMFI) of the CTV label (**Figure 3E**).

**Figure 3.**
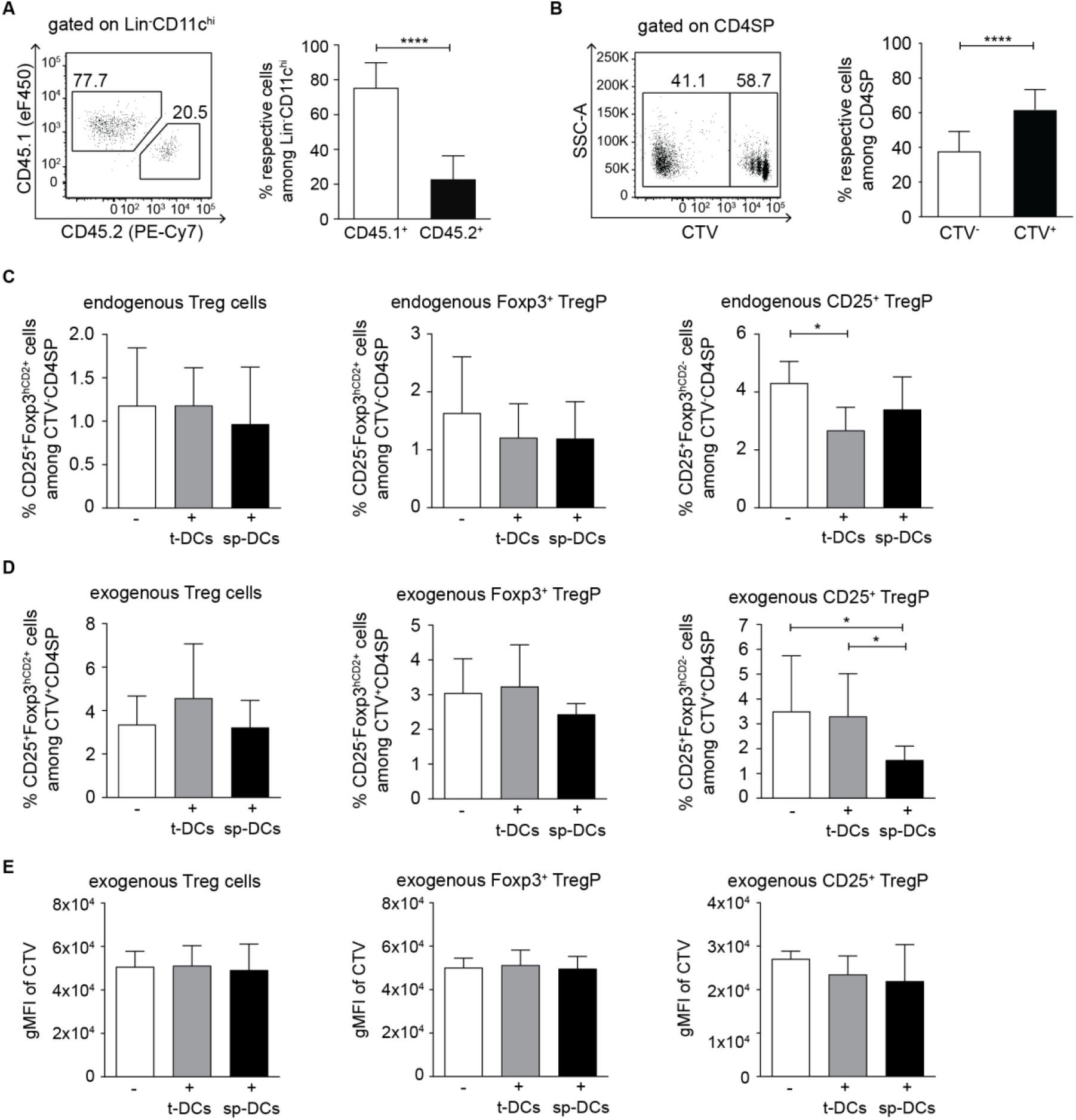
Treg cell frequencies in RTOCs are not affected by introduced sp-DCs. RTOCs were generated from single-cell suspensions of pooled thymi isolated from E14.5 to E16.5 fetuses of Foxp3^hCD2^ reporter mice (BALB/c background), and CTV-labeled CD4SP Foxp3^-^ thymocytes sorted from 4-6 weeks old male Foxp3^hCD2^ reporter mice (BALB/c background) as well as sp-DCs or t-DCs sorted from 4-8 weeks old male CD45.1xBALB/c mice were introduced. **(A)** Introduced cDCs were identified by the congenic marker CD45.1 in the total cDC pool within an RTOC. Representative dot plot (left) and summarizing graph (right) depict frequencies of exogenous (CD45.1^+^) and endogenous (CD45.2^+^) cDCs among Lin^-^CD11c^hi^ cells, respectively. For the representative dot plot, numbers indicate the frequencies of cells within the depicted gates. For the summarizing graph, data are summarized from two independent experiments (mean ± SD; n = 8-15 biological replicates per experiment per group; n = 23 biological replicates per group in total) and tested for significance using Mann-Whitney test; ****p < 0.0001. **(B)** Exogenously added CD4SP thymocytes can be accurately distinguished from endogenous CD4SP thymocytes with the help of the CTV label. Representative dot plot (left) and summarizing graph (right) depict the frequency of CTV^+^ and CTV^-^ cells among CD4SP thymocytes within RTOCs analyzed by flow cytometry on day 4. For the representative dot plot, numbers indicate the frequencies of cells within the depicted gates. For the summarizing graph, data was pooled from RTOCs that contain only CTV-labeled CD4SP thymocytes and RTOCs additionally containing congenically marked t-DCs or sp-DCs from four independent experiments (mean ± SD; n = 7-9 biological replicates per experiment per group; n = 32 biological replicates per group in total) and tested for significance using Mann-Whitney test; ****p < 0.0001. **(C-E)** CTV-labeled CD4SP thymocytes alone or together with sp-DCs and t-DCs, respectively, were introduced into an RTOC. On day 4 of culture, the frequency of CD25^+^Foxp3^hCD2+^ Treg cells, CD25^-^Foxp3^hCD2+^ Treg cell precursors (Foxp3^+^ TregP), and CD25^+^Foxp3^hCD2-^ Treg cell precursors (CD25^+^ TregP) among CTV^-^ (RTOC endogenous) **(C)** and CTV^+^ (RTOC exogenous) **(D)** CD4SP thymocytes was analyzed by flow cytometry. **(E)** On day 4 after set-up, the gMFI of CTV among CD25^+^Foxp3^hCD2+^ Treg cells, CD25^-^Foxp3^hCD2+^ Treg cell precursors (Foxp3^+^ TregP) and CD25^+^Foxp3^hCD2-^ Treg cell precursors (CD25^+^ TregP) among exogenously added CD4SP thymocytes was analyzed by flow cytometry. For **(C-E)**, data are summarized from four independent experiments (mean ± SD; n = 2-3 biological replicates per experiment per group; n = 10-12 biological replicates per group in total). For **(A-E)**, data were tested for significance using Kruskal-Wallis test. Significant differences were indicated by * p<0.05.

Taken together, our data indicate that, although the introduced sp-DCs constitute the majority of the total CD11c^hi^ cDC population within the RTOCs, they did not negatively affect the intrathymic Treg cell induction. This finding suggests that the thymic microenvironment can gradually modulate the phenotype and functional properties of thymus-homing peripheral cDCs.

### The thymic microenvironment modulates the transcriptome of sp-DCs

Next, we assessed how the thymic microenvironment is modulating sp-DCs on a molecular level. To this end, we set up RTOCs and introduced congenically marked *ex vivo* isolated CD45.1^+^Lin^-^CD11c^hi^ sp-DCs (**Figure S4A** in Supplementary Material). RTOCs with introduced CD45.1^+^Lin^-^CD11c^hi^ t-DCs served as controls. At day 2, RTOCs were harvested and CD45.1^+^ sp-DCs or t-DCs were re-isolated (**Figure S4** in Supplementary Material). In this setting, the earlier time point was chosen to study an immediate and direct impact of the thymic microenvironment on the sp-DC phenotype. Low-input RNAseq was performed with these re-isolated cells. As additional controls *ex vivo* isolated CD45^+^Lin^-^CD11c^hi^ sp-DCs and t-DCs were transcriptionally profiled. Comparing sp-DCs re-isolated from RTOCs with *ex vivo* isolated sp-DCs revealed a large number (4262) of differentially expressed genes (DEGs, **Figure 4A**), suggesting that the thymic microenvironment has a noticeable impact on the transcriptome of sp-DCs. Yet, we cannot exclude that the experimental design itself or the rather low sample number also impacted the differential gene expression. Indeed, a distinct fraction of these DEGs, namely 995 genes, were merely impacted by the RTOC itself (‘RTOC effect’, **Figure S5** in Supplementary Material). In contrast, genes assigned as cDC signature genes^25-27^ were not affected by the RTOC microenvironment and equally expressed in all examined groups (**Figure 4B**), while a small number of genes maintained to be differentially expressed between sp-DCs and t-DCs even under RTOC conditions (**Figure 4C**). Among the genes primarily affected in their expression by the ‘RTOC effect’ we found for examples genes involved in cross-presentation (e.g. *Clec4a2, Fcer1g, Fcgr3*), cell proliferation (e.g. *Ccnt1, Ccny, Cdk4, Spag5*) or DC maturation (e.g. *Cd200, Cd274, Relb*) (**Figure 4D**). Interestingly, while genes related to cross-presentation and cell proliferation were mainly negatively impacted by the RTOC, the expression of genes involved in DC maturation, with the exception of MHCII genes, were mainly increased. Importantly, the subset composition of sp-DCs was found to be unaltered within the RTOC compared to input cells when analyzed by flow cytometry (**Figure 4E** and **Figure S6** in Supplementary Material). Thus, although obvious limitations of the culture system exist, the RTOC studies also suggest that the commitment to the cDC1 or cDC2 lineage is a fixed characteristic, probably established already at pre-DC stage.^12, 28^

**Figure 4.**
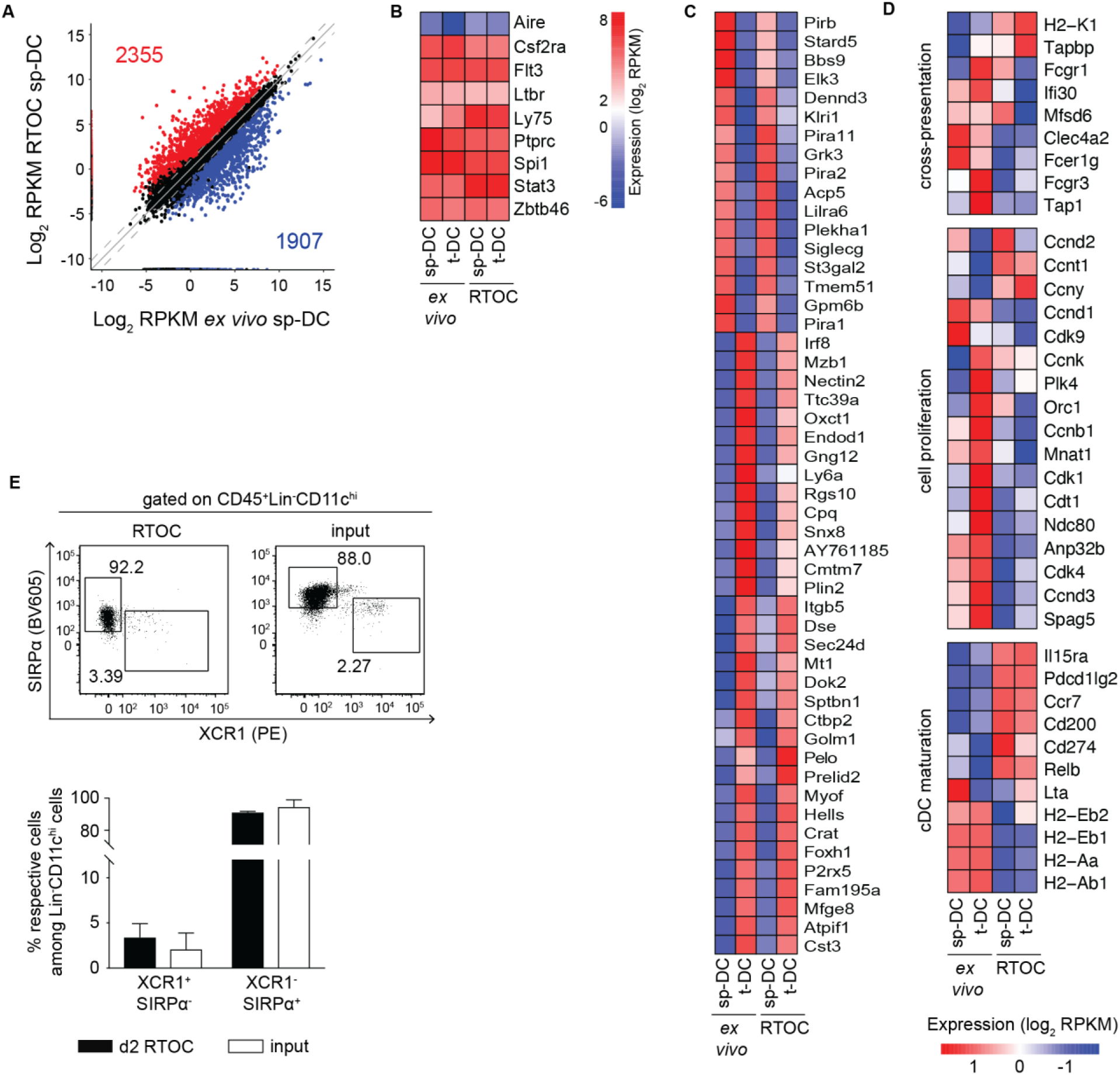
Differences between sp-DCs isolated *ex vivo* or re-isolated from RTOCs are observed on a transcriptomic level. RTOCs were generated from single-cell suspensions of pooled thymi isolated from E14.5 to E16.5 fetuses of Foxp3^hCD2^ reporter mice (BALB/c background), and sp-DCs or t-DCs sorted from 4-8 weeks old female CD45.1xBALB/c mice were introduced. After two days, 1-2×10^3^ cDCs were re-isolated from RTOCs as CD45.1^+^Lin^-^CD11c^hi^ cells by FACS, and total RNA for transcriptional profiling by low-input RNAseq was isolated. Total RNA from cDCs isolated *ex vivo* from 4-8 weeks old female CD45.1xBALB/c mice was used as control. **(A)** Scatter plot highlighting the DEGs between *ex vivo* sp-DCs and RTOC sp-DCs. The DEGs were filtered with a conservative absolute log_2_ fold change (FC) cut-off of at least 1.0 and a p-value cut-off, corrected for multiple testing, of at most 0.05. The RPKM (reads per kilobase of transcript length per million mapped reads) values were calculated based on the variance among samples. **(B)** Heatmap analysis depicting genes for “cross-presentation”, “cell proliferation, and “cDC activation” influenced by the RTOC. Gene sets were manually curated. Bars are color-coded according to the expression value RPKM as indicated in expression scale. Data was mean-centered and rows were clustered using ward.D2 clustering method. **(C)** Heatmap analysis for cDC signature genes that are not differentially expressed between all four cell populations investigated. Bars are color-coded according to the expression value RPKM as indicated in expression scale. **(D)** Heatmap analysis for genes differentially expressed between sp-DCs and t-DCs, independent from *ex vivo* isolation or re-isolation from RTOC. Bars are color-coded according to the expression value RPKM as indicated in expression scale. Columns are clustered by euclidean distances. Data was mean-centered, and rows were clustered using ward.D2 clustering method. **(B-D)** All heatmap analyses were performed on log_2_ transformed data. Each column represents an average of two to three biological replicates. **(E)** The expression of XCR1 and SIRPα on Lin^-^CD11c^hi^ sp-DCs, either sorted from *ex vivo* cells (input) or re-isolated from day 2 RTOCs, was analyzed by flow cytometry. Representative dot plots (top) and summarizing graph (bottom) depict the frequency of XCR1^+^SIRPα^-^ cDC1s and XCR1^-^SIRPα ^+^ cDC2s among Lin^-^CD11c^hi^ sp-DCs. For the representative dot plots, numbers indicate the frequencies of cells within the depicted gates. For the summarizing graph, data are summarized from three independent experiments (mean ± SD; n = 2-8 biological replicates per experiment per group), and data of each subset were independently tested for significance using Mann-Whitney test.

In line with our previously published finding,^16^ we also observed a large number of 2514 DEGs when comparing *ex vivo* isolated sp-DCs and t-DCs (**Figure 5A**). Interestingly, only 857 DEGs were observed between sp-DCs and t-DCs re-isolated from RTOCs, implicating that the thymic microenvironment in RTOCs has a noticeable impact on the transcriptome of sp-DCs, markedly reduced the transcriptional differences between t-DCs and sp-DCs, and finally conferred a transcriptomic profile that more closely resembled the one from t-DCs. In order to identify the genes that were selectively modulated in sp-DCs by the thymic microenvironment of the RTOC, we first determined the overlap of the ‘modulated sp-DC signature’ (i.e., genes likewise up- or down-regulated within sp-DCs re-isolated from RTOCs and *ex vivo* isolated t-DCs when compared to *ex vivo* isolated sp-DCs) with the so called ‘t-DC core signature’, genes that were not differentially regulated between *ex vivo* isolated t-DCs and t-DCs re-isolated from RTOCs (**Figure S7A** in Supplementary Material). Subsequently, from this overlap, all genes that were merely modulated by the RTOC condition itself, the so-called ‘RTOC effect’, were eliminated. This ‘RTOC effect’ was defined as the overlap between three groups of genes: (1) genes commonly expressed in sp-DCs and t-DCs re-isolated from RTOCs, (2) genes differentially regulated between *ex vivo* isolated t-DCs and t-DCs re-isolated from RTOC, and (3) genes differentially regulated between *ex vivo* isolated sp-DCs and sp-DCs re-isolated from RTOC (**Figure S5** in Supplementary Material). By this strict filtering process, 225 genes were identified that were either induced (106) or repressed (119) in sp-DCs by the thymic microenvironment (**Figure 5B** and **Figure S7B** in Supplementary Material). Many of these genes encode immunologically relevant molecules, including “cytokines, chemokines and their respective receptors” (e.g. *Ccl17, Ccl22, Il1r1, Il1a*), “antigen processing associated molecules” (e.g. *Ctsd, Ctsl, Serpinb2, Serpinb10*), “cell adhesion associated molecules” (e.g. *Nedd9, Cd9, Itga5*), “cell migration associated molecules” (e.g. *Cd38, Elmo1, Tubb2b*) and “co-stimulatory molecules” (e.g. *Tnfsf9, Tnfsf4, Cd40, Cd70* and *Cd86*) (**Figure 5C**). Together, these results indicate that the thymic microenvironment can gradually modulate thymus-homing peripheral cDCs on the molecular level by driving the transcriptomic profile of sp-DCs towards the one of t-DCs, thereby transferring thymus-specific properties to the newly entering cDCs from the periphery.

**Figure 5.**
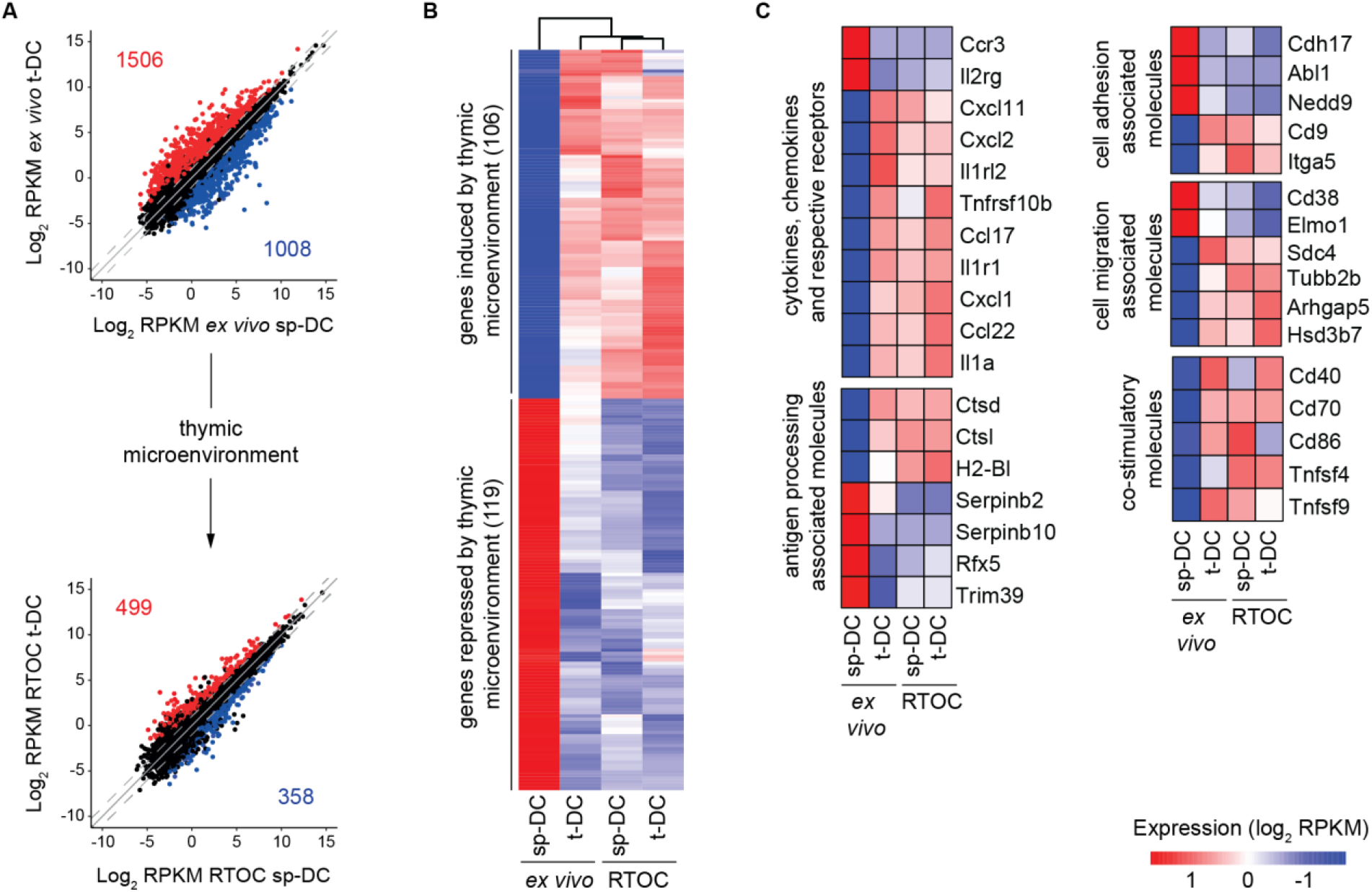
The thymic microenvironment of RTOCs alters the transcriptome of sp-DCs. RTOCs were generated from single-cell suspensions of pooled thymi isolated from E14.5 to E16.5 fetuses of Foxp3^hCD2^ reporter mice (BALB/c background), and sp-DCs or t-DCs sorted from 4-8 weeks old female CD45.1xBALB/c mice were introduced. After two days, 1-2×10^3^ cDCs were re-isolated from RTOCs as CD45.1^+^Lin^-^CD11c^hi^ cells by FACS, and total RNA for transcriptional profiling by low-input RNAseq was isolated. Total RNA from cDCs isolated *ex vivo* from 4-8 weeks old female CD45.1xBALB/c mice was used as control. **(A)** Scatter plots highlighting the DEGs between *ex vivo* cDCs and RTOC cDCs, respectively. The DEGs were filtered with a conservative absolute log_2_ fold change (FC) cut-off of at least 1.0 and a p-value cut-off, corrected for multiple testing, of at most 0.05. The RPKM (reads per kilobase of transcript length per million mapped reads) values were calculated based on the variance among samples. **(B)** A total of 225 genes are shaped by the thymic microenvironment. Heatmap analysis for genes induced or repressed by the thymic microenvironment. Columns are clustered by euclidean distances. **(C)** Heatmap analysis depicting genes for “cytokines, chemokines and their respective receptors”, “antigen processing associated molecules”, “cell adhesion associated molecules”, “cell migration associated molecules”, and “co-stimulatory molecules” influenced by the thymic microenvironment. Gene sets were manually curated. Data was mean-centered and rows were clustered using ward.D2 clustering method. **(B+C)** Heatmaps are color-coded according to the expression value RPKM as indicated in expression scale. All heatmap analyses were performed on log_2_ transformed data. Each column represents an average of two to three biological replicates.

## Discussion

Thymic cDCs play an essential role in key processes of central tolerance like the clonal deletion of self-reactive thymocytes and the generation of Treg cells.^2, 4^ This important contribution to central tolerance was demonstrated for both thymus-resident CD8α^+^SIRPα^-^ t-DCs, which develop intrathymically,^8, 9^ as well as for migratory CD8α^-^SIRPα^+^ t-DCs, which home to the thymus from the periphery via CCR2-mediated chemotaxis and α_4_ integrin-dependent adhesion.^13, 18, 29^ Accordingly, we have previously demonstrated that bulk t-DCs possess a superior Treg cell-inducing capacity when compared to sp-DCs, leading to the differentiation of stable Foxp3^+^ Treg cells.^16^ This raised the question whether sp-DCs, which already have undergone a number of differentiation steps,^30, 31^ are still plastic and can be modulated by the local microenvironment after entry into the thymus to acquire the unique functional properties of t-DCs, including the superior Treg cell-inducing capacity. This point is particularly relevant as several studies have reported diverging transcriptional profiles of cDCs from different lymphoid organs,^32, 33^ suggesting that milieu-specific programs manifest in these different cDCs.

In the present study, we have further addressed this important question experimentally by employing RTOCs to accurately introduce sp-DCs into the thymic microenvironment. For this purpose, RTOCs were carefully chracteriezd regarding their cellular composition, confirming that they mature over time and sucessfully support thymocyte and TEC development. Importantly, we further only used *ex vivo* isolated cDCs from unmanipulated mice, while previously published studies have only investigated the thymus-homing and subsequent maturation of circulating cDCs by adoptively transferring sp-DCs derived from donor mice that were exposed to Flt3L-secreting B16 melanoma cells to expand the cDC population.^17, 29^ Our results suggest that sp-DCs can acquire a thymus-specific functionality, as the addition of an unmanipulated sp-DC population with a physiological subset composition does not negatively impact the Treg cell frequency, although sp-DCs were shown to commonly possess only a low Treg cell-inducing capacity and although these introduced cells constituted the major proportion of total cDCs within the RTOC. These indirect findings suggest that sp-DCs can be modulated by the thymic microenvironment and acquire an improved Treg cell-inducing capacity. Otherwise, the Treg cell frequencies within RTOCs containing the added sp-DCs should have been severely decrease due to the generally poor Treg cell-inducing capacity of sp-DCs.^16^ Interestingly, our results from the control RTOCs, which harbored exogenous t-DCs, further showed that the introduction of additional t-DCs does not increase the Treg cell frequency. This finding is in line with studies demonstrating that the thymic microenvironment creates a saturable niche for the Treg cell development, which is tightly controlled in a TCR-instructive manner by the availability of IL-2.^34-36^ However, as neither the endogenous cDCs nor any other population of the thymic hematopoietic stromal cells (THSCs), such as macrophages or plasmacytoid DCs (pDCs), were depleted from the RTOC, an impact of those cells on the Treg cell frequency cannot be formally excluded. Yet, a considerable impact of any of these populations seems unlikely, because cDCs were shown to constitute the most efficient THSC population in supporting Treg cell development and exogenously added cDCs made up the vast majority of total cDCs within RTOCs.^37^

The molecular profiling of sp-DCs re-isolated from RTOCs implied that the thymic microenvironment tuned the transcriptome of sp-DCs towards the transcriptome of t-DCs already after two days of culture. Yet, it cannot be exclude that the experimental design itself or the rather low sample number may also impacted the differential gene expression. Of note, cDC signature genes and other genes that are relevant for general biological and cellular processes were not changed by the thymic microenvironment. Additionally, our findings support the notion that the subset composition of cDCs is not influenced by the thymus-specific milieu, and commitment to the cDC1 or cDC2 lineage probably occurs already at pre-DC stage.^12, 28^ Among the genes, which were modulated in sp-DCs by the thymic microenvironment, *Ccl17* and *Ccl22* might be of particular relevance as Proietto *et al*. showed that these two chemokines were expressed at very high levels only by CD8α^-^SIRPα^+^ t-DCs, and the supernatant of CD8α^-^SIRPα^+^ t-DCs efficiently attracted CD4SP thymocytes in trans-well migration assays.^38^ Importantly, CCR4, the receptor for CCL17 and CCL22, is expressed on post-positive selection DP and immature CD4SP thymocytes in the medulla and is known to be required for interactions between medullary cDCs and thymocytes.^39^ Hence, the rapid and strong up-regulation of *Ccl17* and *Ccl22* expression on sp-DCs within the thymic microenvironment might confer a higher potency for the clonal deletion of self-reactive thymocytes and the generation of Treg cells through these modulated sp-DCs.

In addition, several co-stimulatory molecules like *Cd40, Cd70, Tnfsf9, Tnfsf4* and *Cd86*, which are known to be essential for the development of Treg cells,^14, 40-42^ were up-regulated on sp-DC re-isolated from RTOCs. Up-regulation of these co-stimulatory molecules follows the steady-state maturation process of t-DCs, which is induced in CD8α^-^SIRPα^+^ migratory t-DCs only upon thymic entry.^11, 17^ Importantly, only matured cDCs possess the capacity to contribute to key processes of central tolerance.^43^ For the intrathymic maturation and homeostasis of cDCs, mTECs were reported to play an only minor role.^44, 45^ Thus, although it is known that the heterogeneity of TECs is completed only in adulthood,^46, 47^ it is unlikely that the observed clear differences in the TEC compartment between embryonic thymi in RTOCs and *ex vivo* isolated adult thymi have a considerable effect on the modulation of sp-DCs. By contrast, different groups recently proposed that cognate interactions of antigen-specific CD4SP thymocytes with both resident and migratory t-DCs efficiently support the homeostatic maturation of t-DCs.^17, 45^ However, in our study, the frequency of CD4SP thymocytes was still rather low in RTOCs harvested on day 2, even though frequencies rapidly increased and reached levels comparable to those observed in *ex vivo* isolated adult thymi by day 4. Thus, it is likely that the initial maturation and adaptation processes seen in sp-DCs re-isolated from RTOCs at day 2 might be imparted by additional mechanisms, while the mechanism involving CD4SP thymocytes might account for the progressed maturation of sp-DCs and acquisition of thymus-specific functional properties after four days of culture. Accordingly, Spidale *et al*. and Oh *et al*. did not rule out the contribution of CD8SP and DP thymocytes to the intrathymic maturation of cDCs.^17, 45^ As these cells are already present in RTOCs at day 2 with a frequency comparable to that found in *ex vivo* isolated adult thymi, they might play a role in the abovementioned initial maturation and adaptation processes. This scenario seems especially likely for DP thymocytes, as they express the chemokine receptor CCR4, like CD4SP thymocytes, and are thus attracted by CCL17 and CCL22,^39^ which we found to be highly induced in sp-DCs re-isolated from RTOCs.

Viewed as a whole, the data from the present study indicate that the thymic microenvironment can moderately modulate the phenotype of sp-DCs, which is likely accompanied by an adjusted Treg cell-inducing capacity. Thus, besides their already advanced differentiation stage cDCs from secondary lymphoid organs actually retain remarkable plasticity, which allows for modulation by the thymic microenvironment. Future studies are required to unravel the exact molecular mechanisms and the precise timing of the intrathymic modulation of thymus-homing peripheral cDCs.

## Material and Methods

### Mice

C.SJL(B6)-*Ptprc*^*a*^*Ptprc*^*b*^/BoyJ mice (CD45.1 congenic mice on BALB/c background, CD45.1xBALB/c mice) and C.Foxp3^tm1(CD2/CD52)Shori^ mice (Foxp3^hCD2^ reporter mice on BALB/c background^48^) were bred and maintained at the central animal facility of the Helmholtz Centre for Infection Research (HZI, Braunschweig, Germany), which provides state-of-the-art laboratory animal care and service. All mice were housed in barriers under specific pathogen-free (SPF) conditions in isolated, ventilated cages, and handled by personnel appropriately trained as well as dedicated animal care staff to assure the highest possible hygienic standards and animal welfare in compliance with German and European animal welfare guidelines. According to the German Animal Welfare Act sacrificing animals solely to remove organs for scientific purposes is notified to the competent authority. This study was carried out in accordance with recommendations defined by FELASA (Federation of European Laboratory Animal Science Associations) and the German animal welfare body GV-SOLAS (Society for Laboratory Animal Science) using approved protocols.

### Antibodies and flow cytometry

Cell suspensions were labeled directly with the following fluorochrome-conjugated anti-mouse antibodies purchased from either BioLegend, BD Biosciences or eBioscience: CD3ε (500A2), CD4 (RM4-5), CD8α (53-6.7), CD11c (N418), CD19 (6D5), CD25 (PC61.5), CD44 (IM7), CD45 (30-F11), CD45.1 (A20), CD45.2 (104), CD49b (DX5), CD80 (16-10A1), CD90.2 (53-2.1), CD172α (P84), CD326 (G8.8), anti-human CD2 (RPA-2.10), Ly51 (6C3), F4/80 (BM8), I-A/I-E (M5/114.15.2), and XCR1 (ZET). UEA-1 was labeled with a biotinylated anti-mouse antibody (clone U1216, Vector Labs) and subsequently detected with a fluorochrome-conjugated streptavidin. To block Fc receptors, the staining mix always contained unconjugated anti-FcRγIII/II antibody (BioXcell; final concentration 10 µg ml^-1^). For exclusion of dead cells, either 4′,6-Diamidine-2′-phenylindole dihydrochloride (Merck) or LIVE/DEAD™ Fixable Near-IR stain kit (Invitrogen) was used. Stained cells were assessed by LSRFortessa™ flow cytometer (BD Biosciences) and data was analyzed with FlowJo® software (TreeStar).

### Isolation of CD4SP Foxp3- thymocytes

CD4^+^CD8^-^ single-positive (SP) Foxp3^-^ thymocytes were isolated from 4-6 weeks old male Foxp3^hCD2^ reporter mice (BALB/c background). Single-cell suspensions from thymi were labeled with anti-hCD2-FITC-conjugated antibody, anti-CD4-PacificBlue-conjugated antibody, anti-CD8α-APC-conjugated antibody and anti-APC microbeads (Miltenyi Biotec). Using the autoMACS® Pro separation system (Miltenyi Biotec), APC-labeled CD8α^+^ cells were depleted. From the negative fraction CD4SP Foxp3^hCD2-^ thymocytes were sorted using a FACSAria™ II (BD Biosciences), a FACSAria™ (BD Biosciences) or a MoFlo XDP (Beckman Coulter).

### Isolation of DCs

Sp-DCs and t-DCs were isolated from 4-8 weeks old CD45.1xBALB/c mice. For low-input RNAseq experiments, female CD45.1xBALB/c mice were used for sp-DC and t-DC isolation. In all other experiments, male CD45.1xBALB/c mice were used for sp-DC and t-DC isolation. After removal of connective tissue, organs were disrupted with the help of scalpels. The tissue fragments were digested in prewarmed Roswell Park Memorial Institute medium (RPMI, Life Technologies) completed with 10 % FCS, 50 U ml^-1^ penicillin, 50 U ml^-1^ streptomycin, 25 mM HEPES, 1 mM sodium pyruvate (all Biochrom AG), and 50 μM β-mercaptoethanol (Life Technologies) (cRPMI), containing 2 mg ml^-1^ collagenase/ dispase (Roche) and 0.2 mg ml^-1^ DNase I (Roche) at 37 °C for 40 min. Digests were filtered through a nylon mesh with pore size of 100 μm (Greiner Bio-One International GmbH) and subjected to a density gradient using high-density Easycoll (1.115 g ml^-1^, Biochrom AG) and low-density Easycoll (1.06 g ml^-1^, Biochrom AG). The gradient was centrifuged at 1350 g, 4 °C with low acceleration and deceleration for 30 min. The low-density interface was collected and cells were stained with respective fluorochrome-conjugated antibodies. Finally, t-DCs were sorted as CD49b^-^F4/80^-^CD90.2^-^CD45.1^+^CD11c^hi^ cells and sp-DCs were sorted as CD49b^-^F4/80^-^CD3^-^CD19^-^CD45.1^+^CD11c^hi^ cells using a FACSAria™ II (BD Biosciences) or a FACSAria™ (BD Biosciences).

### RTOCs

In order to prepare RTOCs, thymi were isolated from E14.5 to E16.5 fetuses of Foxp3^hCD2^ reporter mice (BALB/c background), pooled and digested in cRPMI containing 0.75 mg ml^-1^ collagenase/dispase (Roche) at 37 °C for 35 min. Digestion was stopped by addition of excess medium, and the cell suspension was filtered through a nylon mesh with pore size of 100 μm. 0.65×10^6^ cells were pelleted by centrifugation in a 1.5 ml Eppendorf tube. After complete removal of the supernatant, the cell pellet was dispersed into a slurry and was finally deposited as a freestanding drop on a membrane using a 0.5-2.5 µl Eppendorf pipette. RTOCs were cultured on Whatman® Nuclepore Track-Etched Membrane (0.8 µm pore size, 13 mm diameter; Merck) placed on a sterilized foam sponge (5 mm thick) in one well of a 6-well plate (Sarstedt) containing 4 ml DMEM supplemented with 50 µg ml^-1^ penicillin, 50 µg ml^-1^ streptomycin, 50 µM β-mercaptoethanol, 1 mM non-essential amino acids (all from Life Technologies), 10 % FCS and 10 mM HEPES (both from Biochrom AG). Culture conditions were 37 °C and 5 % CO_2_.

For phenotypic characterization, RTOCs were collected with forceps and 2-3 pooled RTOCs were prepared for analysis on day 2 and day 4. RTOCs analyzed for the composition of the T cell compartment were mechanically disrupted with the help of a plunger and a nylon mesh with pore size of 100 μm. For the analysis of the composition of the TEC compartment, RTOCs were digested in cRPMI (Life Technologies) supplemented with 1 mg ml^-1^ collagenase/dispase (Roche) and 0.1 mg ml^-1^ DNase I (Roche) with gentle rocking at 37 °C for 30 min. To support disaggregation of the RTOCs, the digestion suspension was mixed by pipetting every 5-10 min during incubation.

RTOCs containing cDCs were prepared by addition of 0.1×10^6^ sorted sp-DCs or t-DCs to the 0.65×10^6^ cells from the fetal thymi before pelleting. In order to perform low-input RNAseq on the inserted sp-DCs and t-DCs, RTOCs were harvested on day 2, 2-3 RTOCs were pooled and processed by digestion.

For the set-up of syngenic RTOC co-cultures, 0.65×10^6^ cells from the fetal thymi were mixed with 0.5×10^5^ Cell Trace Violet (CTV)™ (Invitrogen)-labeled sorted CD4SP Foxp3^hCD2-^ thymocytes and 0.1×10^6^ sorted sp-DCs or t-DCs before pelleting. On day 4, RTOCs were collected with forceps, 2-3 RTOCs were pooled and processed by mechanical disruption for flow cytometric analysis.

### Low-input RNAseq

Total RNA from 1-2×10^3^ CD45.1^+^Lin^-^CD11c^hi^ cDCs, which were either isolated directly *ex vivo* or re-isolated from RTOCs, was obtained using the RNeasy® Plus Micro Kit (Qiagen). cDNA was synthesized from 1 ng RNA using the SMART-Seq® v4 Ultra® Low Input RNA Kit for Sequencing (Takara Bio Europe SAS). Libraries were prepared from the purified cDNA using the Nextera™ XT DNA Library Preparation Kit (Illumina). As input, 0.1 ng cDNA was used per sample and the libraries were cleaned up with 1.8x AMPure® XP beads (Beckman Coulter). Quality and integrity of nucleic acids were assessed using the Agilent Technologies 2100 Bioanalyzer (Agilent Technologies) after each step. The generated libraries were sequenced at the Genome Analytics facility of the Helmholtz Centre for Infection Research on an Illumina HiSeq2500 using 50 bp single reads. The sequenced libraries were assessed for read quality with the FastQC tool (http://www.bioinformatics.babraham.ac.uk/projects/fastqc), and alignment of the libraries versus the mouse reference genome (Assembly: GRCm38.p6) was performed using the splice junction mapper TopHat2 v1.2.0 ^49^ with default parameterization. Subsequently, DESeq2 ^50^ was used to compute the differential gene expression between the four different conditions from the read counts. The differentially expressed genes (DEGs) were filtered with a conservative absolute log_2_ fold change (FC) cut-off of at least 1.0 and a p-value cut-off, corrected for multiple testing, of at most 0.05. Additionally, based on the variance among samples, the RPKM (reads per kilobase of transcript length per million mapped reads) values were calculated.

### Statistical analysis

The GraphPad Prism software v7.0 (Graph-Pad) was used to perform all statistical analyses. Data are presented as mean±standard deviation (SD). For comparison of unmatched groups, two-tailed Mann-Whitney test was applied and the p-values were calculated with long-rank test (Mantel-Cox). If comparing more than three groups, Kruskal-Wallis-Test was used. A p-value below 0.05 was considered as significant; * p<0.05; ** p<0.01; *** p<0.001; **** p<0.0001; ns (not significant).

## Acknowledgements

We thank Lothar Groebe, Maria Hoexter and Petra Hagendorff for cell sorting, and the Genome Analytics facility for RNAseq.

## Author Contributions

SH performed the experiments. SH and MB analyzed the data. SH and JH designed the research, interpreted the data, and wrote the manuscript.

## Data availability

RNAseq data can be accessed at NCBI GEO under the accession number GSE164280 (www.ncbi.nlm.nih.gov/geo/query/acc.cgi?acc=GSE164280).

## References

1. Gray DH, Ueno T, Chidgey AP, et al. Controlling the thymic microenvironment. Curr Opin Immunol. 2005;17(2):137–43.

2. Klein L, Kyewski B, Allen PM, Hogquist KA. Positive and negative selection of the T cell repertoire: what thymocytes see (and don’t see). Nat Rev Immunol. 2014;14(6):377–91.

3. Klein L, Robey EA, Hsieh CS. Central CD4^+^ T cell tolerance: deletion versus regulatory T cell differentiation. Nat Rev Immunol. 2019;19(1):7–18.

4. Wirnsberger G, Hinterberger M, Klein L. Regulatory T-cell differentiation versus clonal deletion of autoreactive thymocytes. Immunol Cell Biol. 2011;89(1):45–53.

5. Gurka S, Hartung E, Becker M, Kroczek RA. Mouse Conventional Dendritic Cells Can be Universally Classified Based on the Mutually Exclusive Expression of XCR1 and SIRPα. Front Immunol. 2015;6:35.

6. Wu L, Li CL, Shortman K. Thymic dendritic cell precursors: relationship to the T lymphocyte lineage and phenotype of the dendritic cell progeny. J Exp Med. 1996;184(3):903–11.

7. Lei Y, Ripen AM, Ishimaru N, et al. Aire-dependent production of XCL1 mediates medullary accumulation of thymic dendritic cells and contributes to regulatory T cell development. J Exp Med. 2011;208(2):383–94.

8. Lyszkiewicz M, Zietara N, Fohse L, et al. Limited niche availability suppresses murine intrathymic dendritic-cell development from noncommitted progenitors. Blood. 2015;125(3):457–64.

9. Perry JSA, Lio CJ, Kau AL, et al. Distinct contributions of Aire and antigen-presenting-cell subsets to the generation of self-tolerance in the thymus. Immunity. 2014;41(3):414–26.

10. Wu L, D’Amico A, Hochrein H, O’Keeffe M, Shortman K, Lucas K. Development of thymic and splenic dendritic cell populations from different hemopoietic precursors. Blood. 2001;98(12):3376–82.

11. Li J, Park J, Foss D, Goldschneider I. Thymus-homing peripheral dendritic cells constitute two of the three major subsets of dendritic cells in the steady-state thymus. J Exp Med. 2009;206(3):607–22.

12. Schlitzer A, Sivakamasundari V, Chen J, et al. Identification of cDC1- and cDC2-committed DC progenitors reveals early lineage priming at the common DC progenitor stage in the bone marrow. Nat Immunol. 2015;16(7):718–28.

13. Baba T, Nakamoto Y, Mukaida N. Crucial contribution of thymic Sirpα^+^ conventional dendritic cells to central tolerance against blood-borne antigens in a CCR2-dependent manner. J Immunol. 2009;183(5):3053–63.

14. Coquet JM, Ribot JC, Babala N, et al. Epithelial and dendritic cells in the thymic medulla promote CD4^+^Foxp3^+^ regulatory T cell development via the CD27-CD70 pathway. J Exp Med. 2013;210(4):715–28.

15. Oh J, Wu N, Baravalle G, et al. MARCH1-mediated MHCII ubiquitination promotes dendritic cell selection of natural regulatory T cells. J Exp Med. 2013;210(6):1069–77.

16. Garg G, Nikolouli E, Hardtke-Wolenski M, et al. Unique properties of thymic antigen-presenting cells promote epigenetic imprinting of alloantigen-specific regulatory T cells. Oncotarget. 2017;8(22):35542–57.

17. Oh J, Wu N, Barczak AJ, Barbeau R, Erle DJ, Shin JS. CD40 Mediates Maturation of Thymic Dendritic Cells Driven by Self-Reactive CD4^+^ Thymocytes and Supports Development of Natural Regulatory T Cells. J Immunol. 2018;200(4):1399–412.

18. Proietto AI, van Dommelen S, Zhou P, et al. Dendritic cells in the thymus contribute to T-regulatory cell induction. Proc Natl Acad Sci U S A. 2008;105(50):19869–74.

19. Jenkinson EJ, Anderson G, Owen JJ. Studies on T cell maturation on defined thymic stromal cell populations in vitro. J Exp Med. 1992;176(3):845–53.

20. Rothenberg EV, Moore JE, Yui MA. Launching the T-cell-lineage developmental programme. Nat Rev Immunol. 2008;8(1):9–21.

21. Gill J, Malin M, Sutherland J, Gray D, Hollander G, Boyd R. Thymic generation and regeneration. Immunol Rev. 2003;195:28–50.

22. Anderson G, Jenkinson EJ, Moore NC, Owen JJ. MHC class II-positive epithelium and mesenchyme cells are both required for T-cell development in the thymus. Nature. 1993;362(6415):70–3.

23. Seach N, Wong K, Hammett M, Boyd RL, Chidgey AP. Purified enzymes improve isolation and characterization of the adult thymic epithelium. J Immunol Methods. 2012;385(1-2):23–34.

24. Abramson J, Anderson G. Thymic Epithelial Cells. Annu Rev Immunol. 2017;35:85–118.

25. Merad M, Sathe P, Helft J, Miller J, Mortha A. The dendritic cell lineage: ontogeny and function of dendritic cells and their subsets in the steady state and the inflamed setting. Annu Rev Immunol. 2013;31:563–604.

26. Miller JC, Brown BD, Shay T, et al. Deciphering the transcriptional network of the dendritic cell lineage. Nat Immunol. 2012.

27. Mildner A, Jung S. Development and function of dendritic cell subsets. Immunity. 2014;40(5):642–56.

28. Grajales-Reyes GE, Iwata A, Albring J, et al. Batf3 maintains autoactivation of Irf8 for commitment of a CD8α^+^ conventional DC clonogenic progenitor. Nat Immunol. 2015;16(7):708–17.

29. Bonasio R, Scimone ML, Schaerli P, Grabie N, Lichtman AH, von Andrian UH. Clonal deletion of thymocytes by circulating dendritic cells homing to the thymus. Nat Immunol. 2006;7(10):1092–100.

30. Kamath AT, Henri S, Battye F, Tough DF, Shortman K. Developmental kinetics and lifespan of dendritic cells in mouse lymphoid organs. Blood. 2002;100(5):1734–41.

31. Liu K, Waskow C, Liu X, Yao K, Hoh J, Nussenzweig M. Origin of dendritic cells in peripheral lymphoid organs of mice. Nat Immunol. 2007;8(6):578–83.

32. Elpek KG, Bellemare-Pelletier A, Malhotra D, et al. Lymphoid organ-resident dendritic cells exhibit unique transcriptional fingerprints based on subset and site. PLoS One. 2011;6(8):e23921.

33. Mahiddine K, Hassel C, Murat C, Girard M, Guerder S. Tissue-Specific Factors Differentially Regulate the Expression of Antigen-Processing Enzymes During Dendritic Cell Ontogeny. Front Immunol. 2020;11:453.

34. Bautista JL, Lio CW, Lathrop SK, et al. Intraclonal competition limits the fate determination of regulatory T cells in the thymus. Nat Immunol. 2009;10(6):610–7.

35. Leung MW, Shen S, Lafaille JJ. TCR-dependent differentiation of thymic Foxp3^+^ cells is limited to small clonal sizes. J Exp Med. 2009;206(10):2121–30.

36. Weist BM, Kurd N, Boussier J, Chan SW, Robey EA. Thymic regulatory T cell niche size is dictated by limiting IL-2 from antigen-bearing dendritic cells and feedback competition. Nat Immunol. 2015;16(6):635–41.

37. Guerri L, Peguillet I, Geraldo Y, Nabti S, Premel V, Lantz O. Analysis of APC types involved in CD4 tolerance and regulatory T cell generation using reaggregated thymic organ cultures. J Immunol. 2013;190(5):2102–10.

38. Proietto AI, Lahoud MH, Wu L. Distinct functional capacities of mouse thymic and splenic dendritic cell populations. Immunol Cell Biol. 2008;86(8):700–8.

39. Hu Z, Lancaster JN, Sasiponganan C, Ehrlich LI. CCR4 promotes medullary entry and thymocyte-dendritic cell interactions required for central tolerance. J Exp Med. 2015;212(11):1947–65.

40. Watanabe N, Wang YH, Lee HK, et al. Hassall’s corpuscles instruct dendritic cells to induce CD4^+^CD25^+^ regulatory T cells in human thymus. Nature. 2005;436(7054):1181–5.

41. Mahmud SA, Manlove LS, Schmitz HM, et al. Costimulation via the tumor-necrosis factor receptor superfamily couples TCR signal strength to the thymic differentiation of regulatory T cells. Nat Immunol. 2014;15(5):473–81.

42. Tai X, Cowan M, Feigenbaum L, Singer A. CD28 costimulation of developing thymocytes induces Foxp3 expression and regulatory T cell differentiation independently of interleukin 2. Nat Immunol. 2005;6(2):152–62.

43. Ardouin L, Luche H, Chelbi R, et al. Broad and Largely Concordant Molecular Changes Characterize Tolerogenic and Immunogenic Dendritic Cell Maturation in Thymus and Periphery. Immunity. 2016;45(2):305–18.

44. Cosway EJ, Lucas B, James KD, et al. Redefining thymus medulla specialization for central tolerance. J Exp Med. 2017;214(11):3183–95.

45. Spidale NA, Wang B, Tisch R. Cutting edge: Antigen-specific thymocyte feedback regulates homeostatic thymic conventional dendritic cell maturation. J Immunol. 2014;193(1):21–5.

46. Han J, Zuniga-Pflucker JC. A 2020 View of Thymus Stromal Cells in T Cell Development. J Immunol. 2021;206(2):249–56.

47. Kadouri N, Nevo S, Goldfarb Y, Abramson J. Thymic epithelial cell heterogeneity: TEC by TEC. Nat Rev Immunol. 2020;20(4):239–53.

48. Komatsu N, Mariotti-Ferrandiz ME, Wang Y, Malissen B, Waldmann H, Hori S. Heterogeneity of natural Foxp3^+^ T cells: a committed regulatory T-cell lineage and an uncommitted minor population retaining plasticity. Proc Natl Acad Sci U S A. 2009;106(6):1903–8.

49. Kim D, Pertea G, Trapnell C, Pimentel H, Kelley R, Salzberg SL. TopHat2: accurate alignment of transcriptomes in the presence of insertions, deletions and gene fusions. Genome Biol. 2013;14(4):R36.

50. Love MI, Huber W, Anders S. Moderated estimation of fold change and dispersion for RNA-seq data with DESeq2. Genome Biol. 2014;15(12):550.

