## Supplementary Material for "The thymic microenvironment gradually modulates the phenotype of thymus-homing peripheral conventional dendritic cells"

### Supplementary Figures

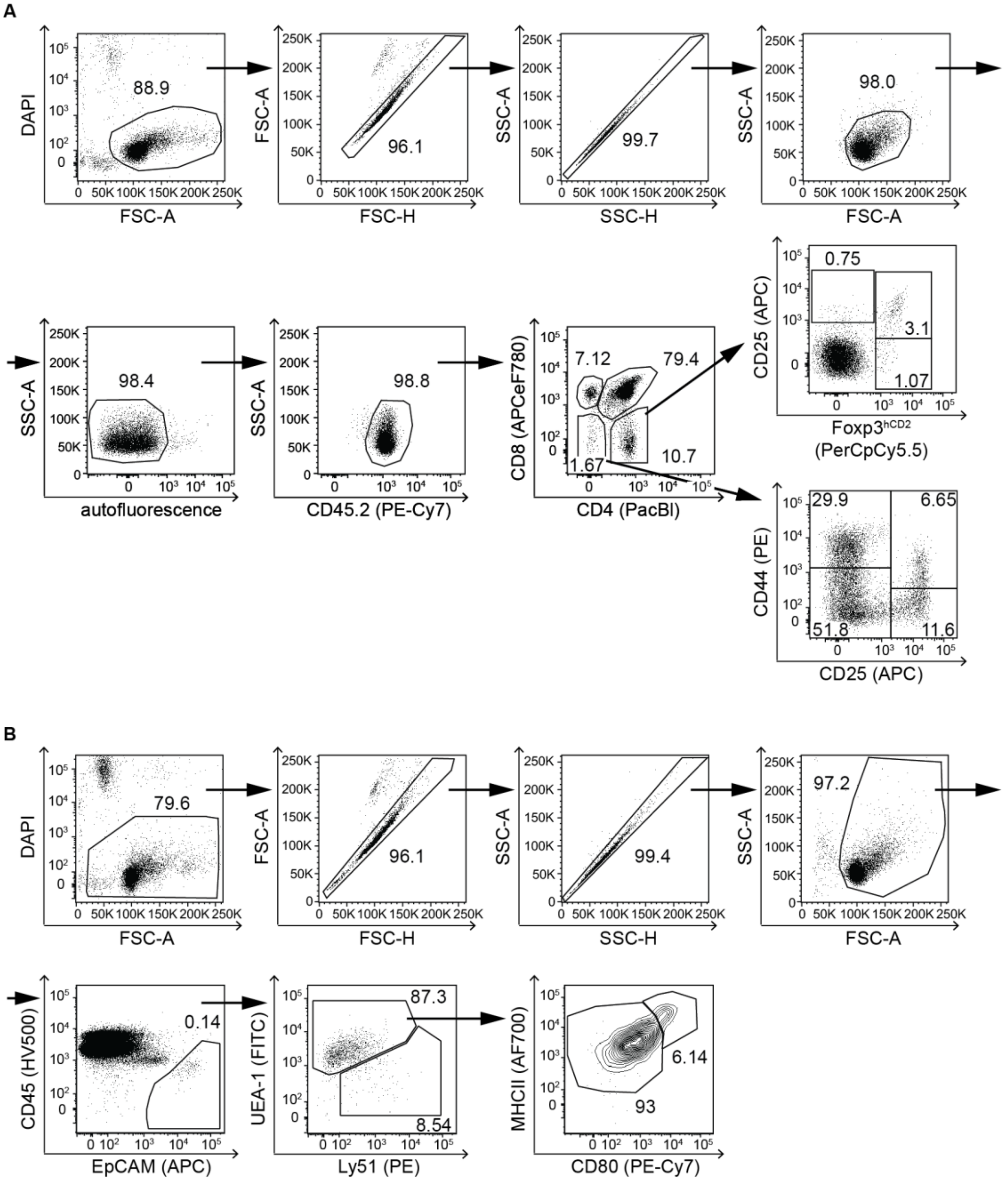


**Figure S1. Phenotypic analysis of the thymocyte and TEC compartment within RTOCs. (A)** Exemplary gating strategy to identify the major thymocyte populations in *ex vivo* isolated thymi and RTOCs by flow cytometry. Numbers indicate the frequencies of cells within the depicted gates. **(B)** Exemplary gating strategy to identify the major TEC populations and subsets in *ex vivo* isolated thymi and RTOCs by flow cytometry. Numbers indicate the frequencies of cells within the depicted gates.

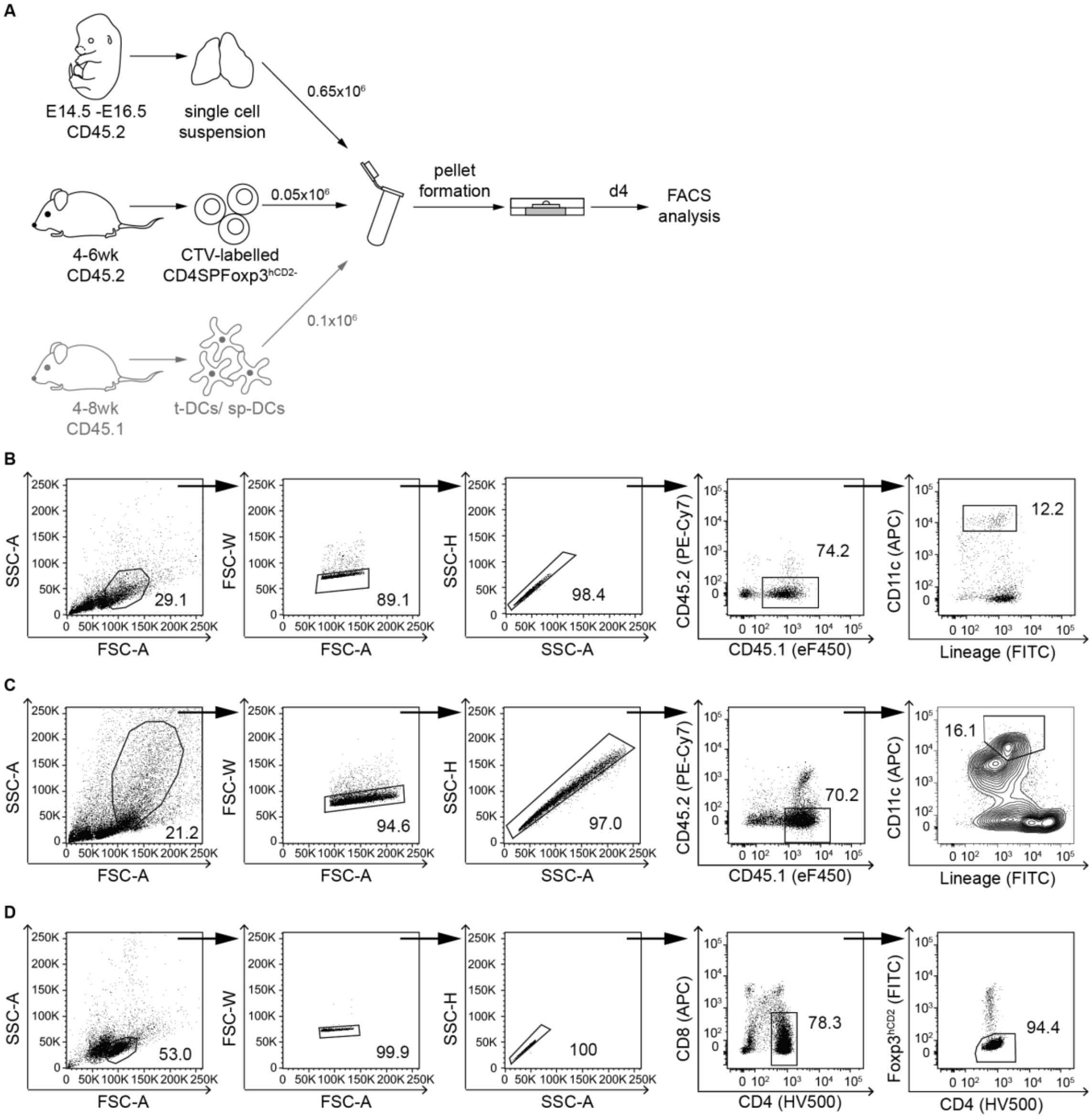


**Figure S2. Set-up of syngenic RTOC co-cultures. (A)** To set up syngenic co-cultures within an RTOC, $Lin^{-}CD11c^{hi}$ t-DCs and sp-DCs isolated from 4-8 weeks old male CD45.1xBALB/c mice, CTV-labeled CD4SP $Foxp3^{hCD2-}$ cells isolated from 4-6 weeks old male $Foxp3^{hCD2}$ reporter mice (BALB/c, CD45.2), and total single-cell suspensions of pooled thymi isolated from E14.5-E16.5 fetuses of $Foxp3^{hCD2}$ reporter mice (BALB/c, CD45.2) were mixed, pelleted, and analyzed on day 4. **(B)** Exemplary gating strategy to sort sp-DCs from *ex vivo* isolated spleens of 4-8 weeks old male CD45.1xBALB/c mice by fluorescence-activated cell sorting. Numbers indicate the frequencies of cells within the depicted gates. **(C)** Exemplary gating strategy to sort t-DCs from *ex vivo* isolated thymi of 4-8 weeks old male CD45.1xBALB/c mice by fluorescence-activated cell sorting. Numbers indicate the frequencies of cells within the depicted gates. **(D)** Exemplary gating strategy to sort CD4SP $Foxp3^{hCD2-}$ cells from *ex vivo* isolated thymi of 4-6 weeks old male $Foxp3^{hCD2}$ reporter mice (BALB/c, CD45.2) by fluorescence-activated cell sorting. Numbers indicate the frequencies of cells within the depicted gates.

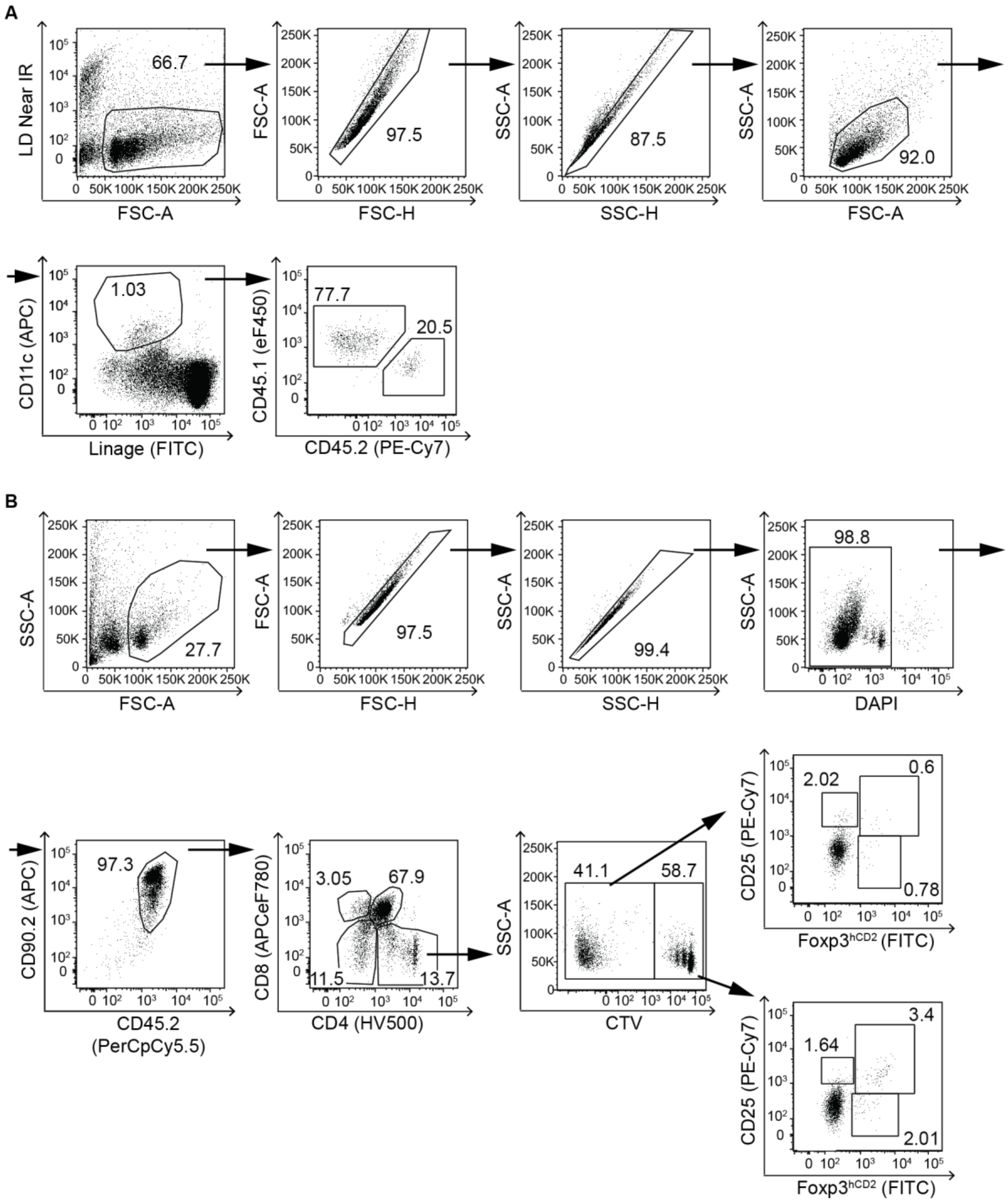


**Figure S3. Analysis of syngenic RTOC co-cultures. (A)** Exemplary gating strategy to identify exogenously added CD45.1$^{+}$ cDCs and their endogenous CD45.2$^{+}$ counterparts by flow cytometry in RTOCs harvested on day 4. Numbers indicate the frequencies of cells within the depicted gates. **(B)** Exemplary gating strategy to identify CTV$^{-}$ (RTOC endogenous) and CTV$^{+}$ (RTOC exogenous) Treg cells by flow cytometry in RTOCs harvested on day 4. Numbers indicate the frequencies of cells within the depicted gates.

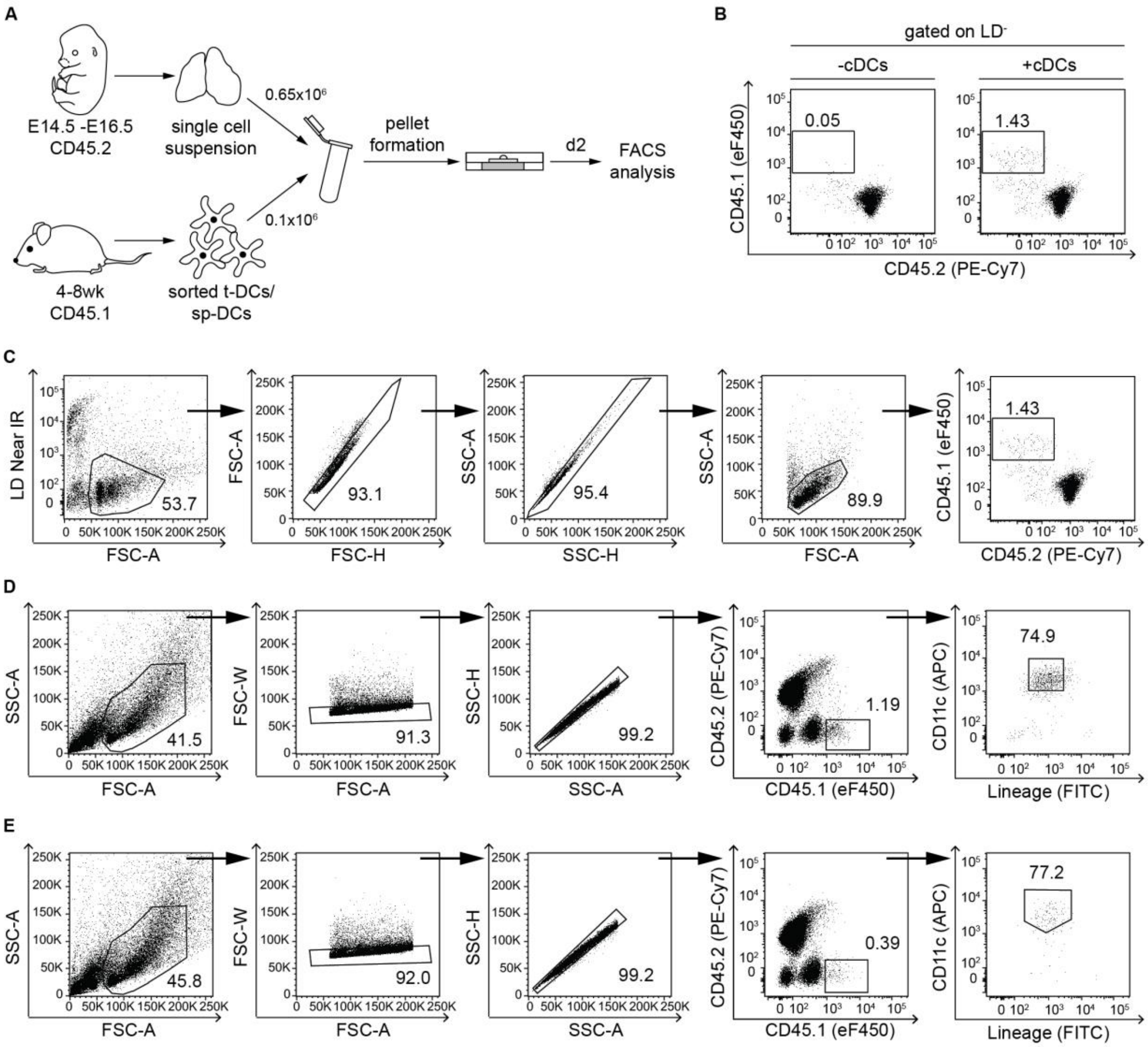


**Figure S4. cDCs can be introduced into RTOCs and re-isolated with the help of congenic markers.** **(A)** 0.1x10$^6$ Lin$^-$CD11c$^{hi}$ t-DCs or Lin$^-$CD11c$^{hi}$ sp-DCs (Lin defined as CD49b, F4/80 and CD90 or CD49b, F4/80, CD3 and CD19) isolated from 4-8 weeks old female CD45.1xBALB/c mice were introduced into an RTOC consisting of 0.65x10$^6$ single cells of pooled thymi isolated from E14.5-E16.5 fetuses of Foxp3$^{hCD2}$ reporter mice (BALB/c, CD45.2) for two days. (**B**) Flow cytometric analysis of an RTOC on day 2 shows that the introduced cDCs can be re-identified by differential congenic markers (right), while RTOCs consisting exclusively of total single-cell suspensions of pooled thymi isolated from E14.5-E16.5 fetuses of Foxp3$^{hCD2}$ reporter mice (BALB/c, CD45.2) do not contain any CD45.1$^+$ cells (left). Representative dot plots for RTOCs containing sp-DCs show frequency of CD45.1$^+$ and CD45.2$^+$ cells among living cells. **(C)** Exemplary gating strategy to identify exogenously added CD45.1$^+$ cDCs by flow cytometry in RTOCs harvested on day 2. Numbers indicate the frequencies of cells within the depicted gates. **(D)** Exemplary gating strategy to re-isolate sp-DCs from RTOCs on day 2 after set-up by fluorescence-activated cell sorting. Numbers indicate the frequencies of cells within the depicted gates. **(E)** Exemplary gating strategy to re-isolate t-DCs from RTOCs on day 2 after set-up by fluorescence-activated cell sorting. Numbers indicate the frequencies of cells within the depicted gates.

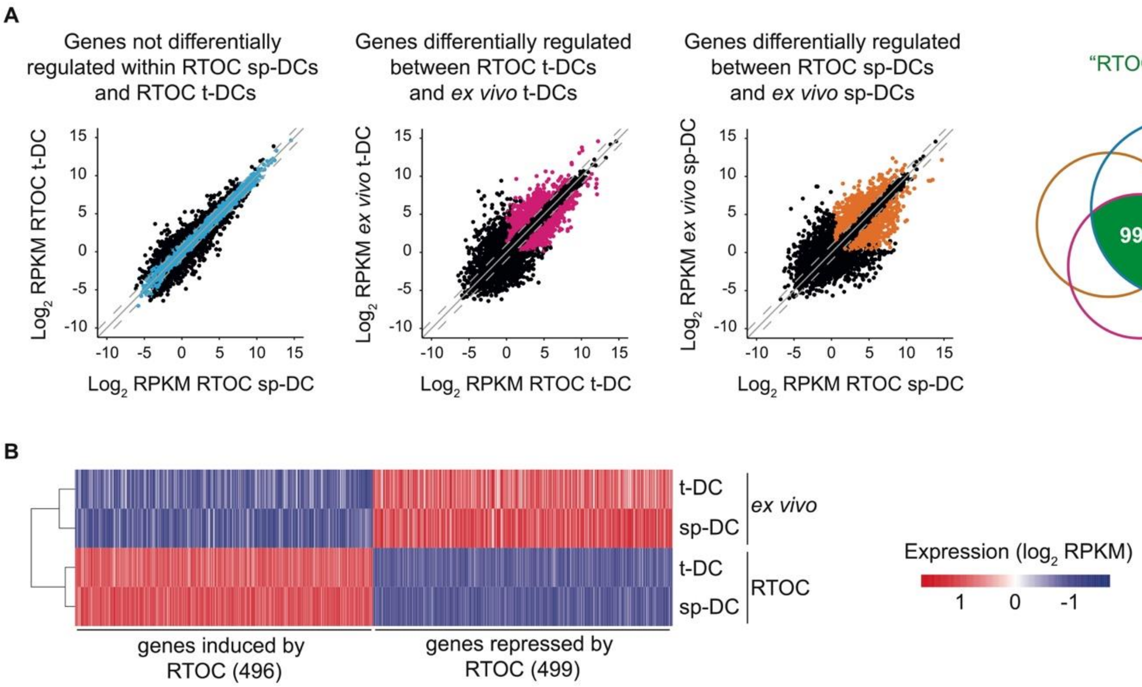


**Figure S5. A total of 995 genes are influenced by the RTOC. (A)** Scatter plots (left) depicting genes not differentially regulated within RTOC sp-DCs and RTOC t-DCs (blue), genes differentially regulated between RTOC t-DCs and *ex vivo* t-DCs (pink) and genes differentially regulated between RTOC sp-DCs and *ex vivo* sp-DCs (orange). Venn diagram (right) depicting the contribution of these three gene sets to the RTOC signature. **(B)** Heatmap analysis for genes induced or repressed by the RTOC. Heatmap analysis was performed on $\log_2$ transformed RPKM. Bars are color-coded according to the expression value RPKM as indicated in expression scale. Data represents the mean of two to three biological replicates per condition. Data was mean-centered, rows were clustered using ward.D2 clustering method and columns clustered based on the Euclidean distance.

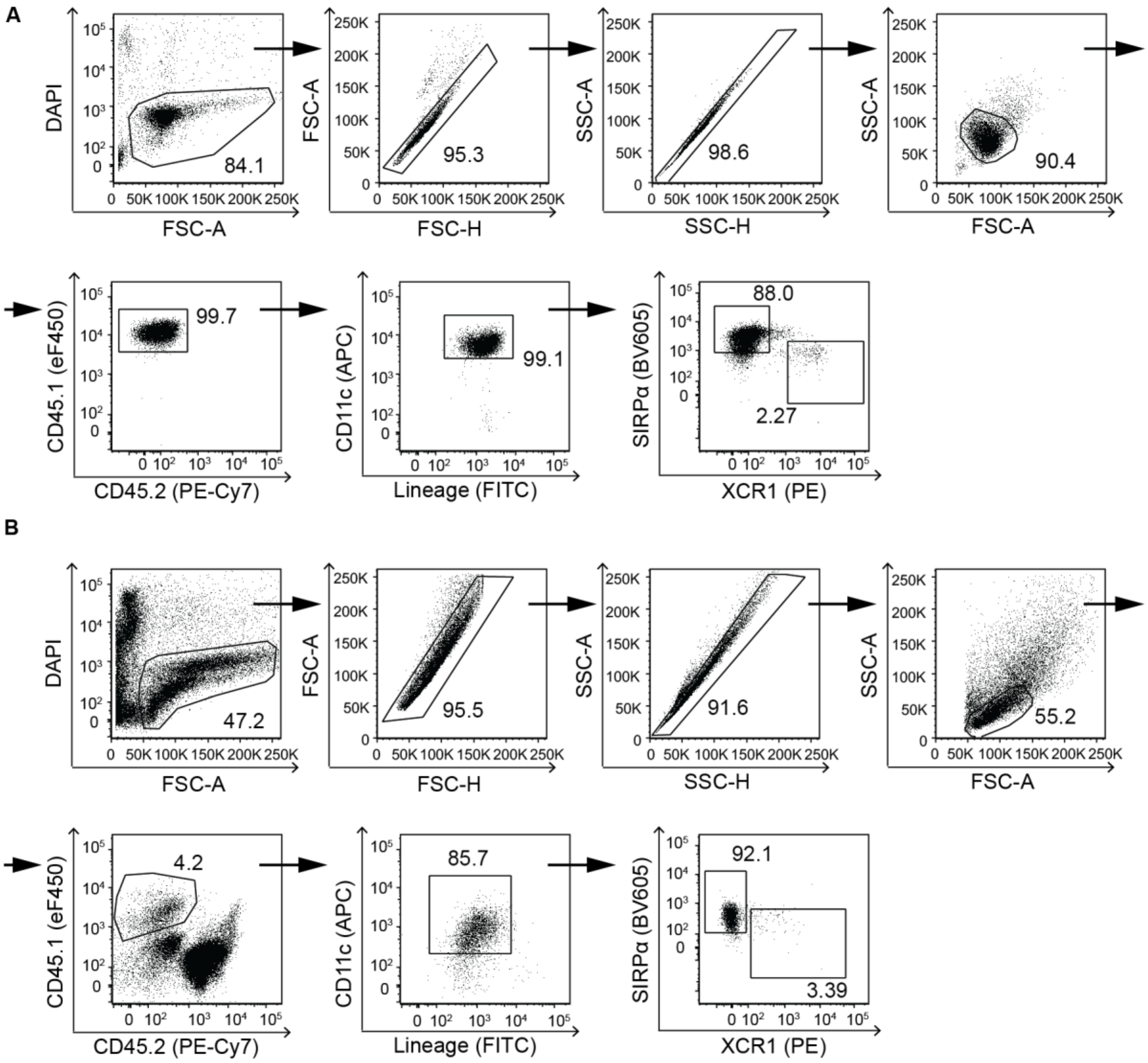


**Figure S6. Subset composition of sp-DCs.** The expression of XCR1 and SIRPα on Lin$^-$CD11c$^{hi}$ sp-DCs, isolated either *ex vivo* or re-isolated from day 2 RTOCs, was analyzed by flow cytometry. Exemplary gating strategy to identify XCR1$^+$SIRPα$^-$ cDC1s and XCR1$^-$SIRPα$^+$ cDC2s among Lin$^-$CD11c$^{hi}$ sp-DCs sorted from *ex vivo* cells (input) **(A)** or re-isolated from RTOCs harvested on day 2 **(B)**.

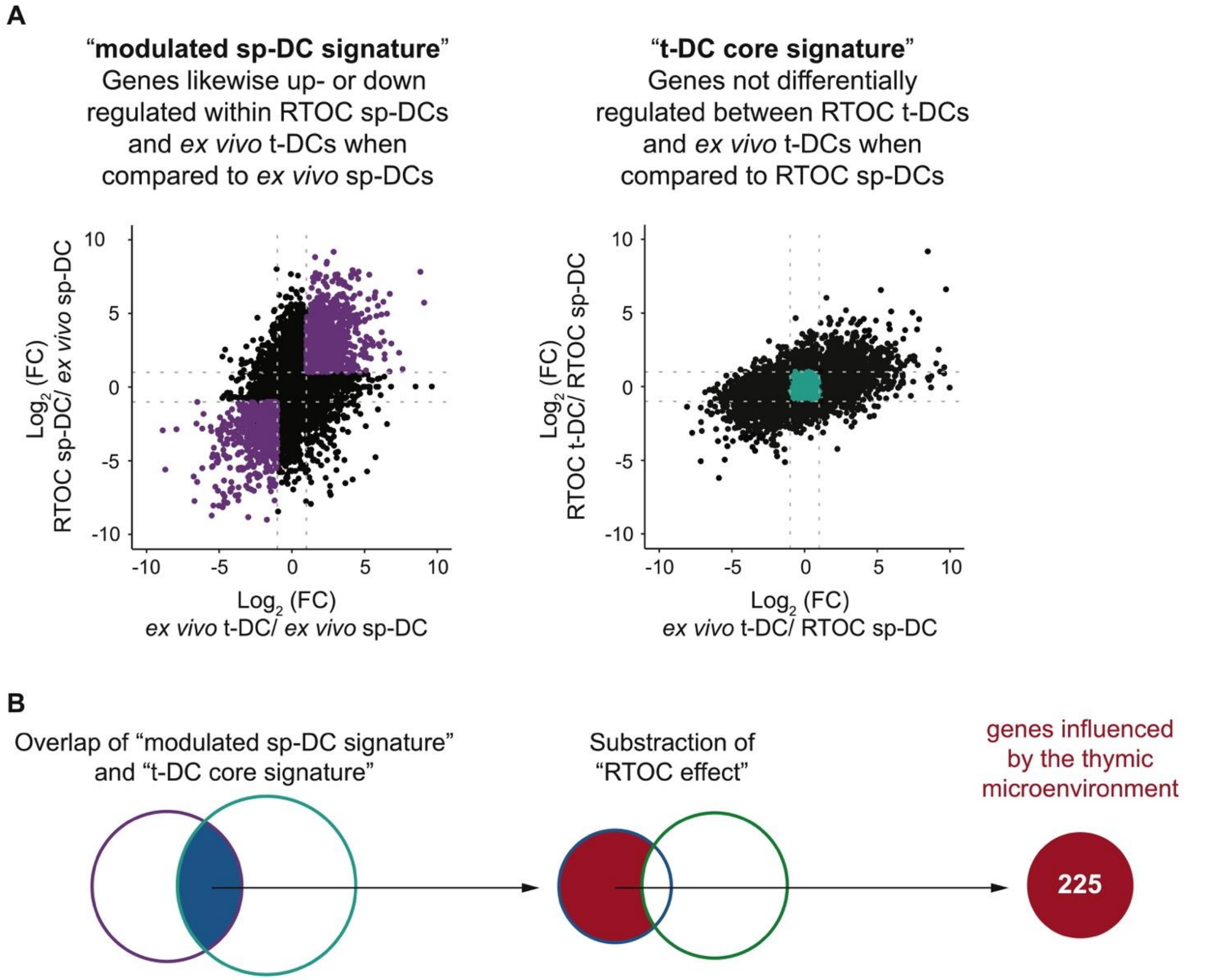


**Figure S7. Filtering strategy applied to reveal cDC genes influenced by the thymic microenvironment. (A)** FC plots illustrating the two overlapping conditions, which are considered to identify genes defined as influenced by the thymic microenvironment. Only genes, which are likewise up- or down regulated within RTOC sp-DCs and *ex vivo* t-DCs when compared to *ex vivo* sp-DCs ('modulated sp-DC signature', purple, left) and at the same time not differentially regulated between RTOC t-DCs and *ex vivo* t-DCs when compared to RTOC sp-DCs ('t-DC core signature', cyan, right) were considered to be influenced in sp-DCs by the thymic microenvironment. **(B)** Venn diagrams exemplifying filtering from all DEGs to the 225 genes that are influenced by the thymic microenvironment. For this purpose, the overlap of 'modulated sp-DC signature' and 't-DC core signature' was subtracted by the 'RTOC-effect'.
